## Supplementary material for "Genomic deletion of Bcl6 differentially affects conventional dendritic cell subsets and compromises Tfh/Tfr/Th17 cell responses"

**<u>Supplementary Figure file</u>**

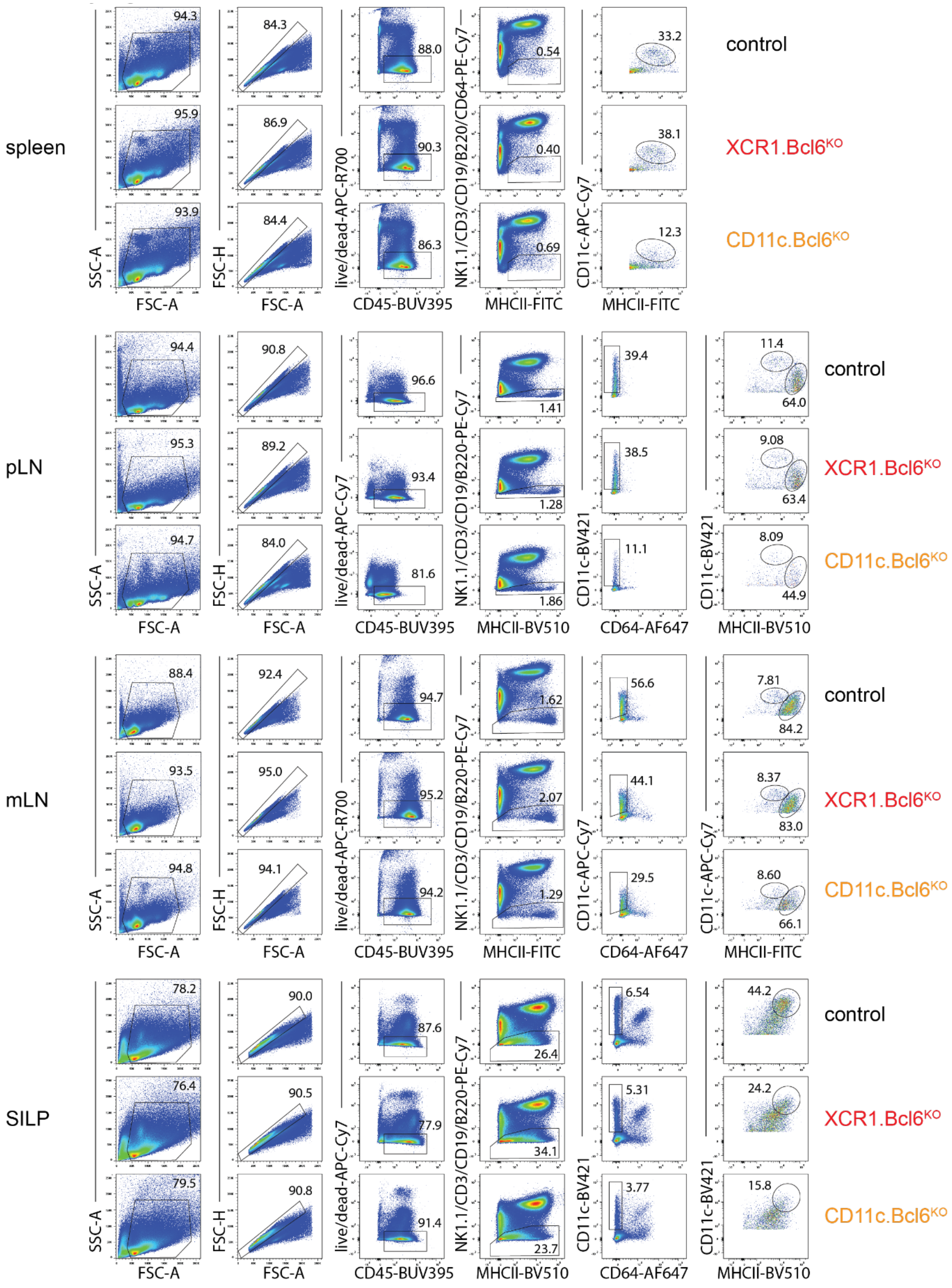


**Supplementary Figure 1:** Full gating strategy for spleen, pLN, mLN and SILP cDC from control, *XCR1.Bcl6*$^{KO}$, and *CD11c.Bcl6*$^{KO}$ mice (used for main article Figures 1 and 2).

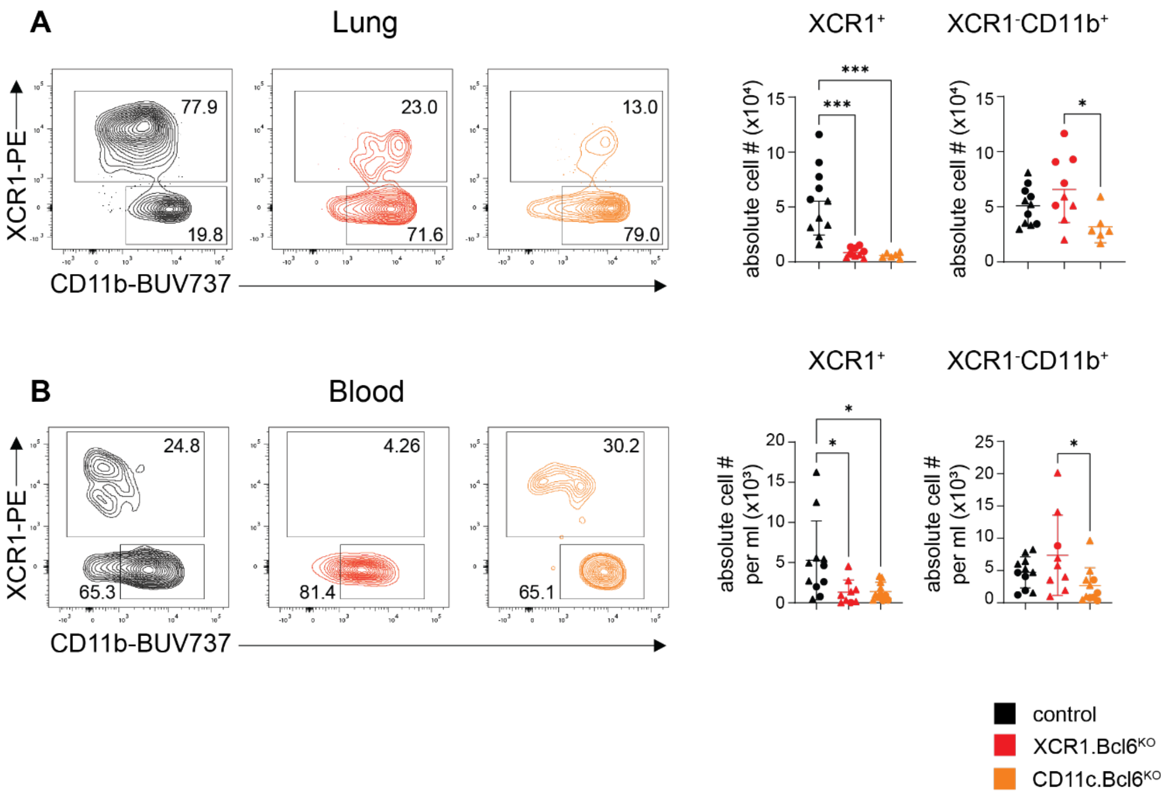


**Supplementary Figure 2**

A. Left: Flow cytometry plots show XCR1⁺ and XCR1⁻CD11b⁺ DC subsets in the lungs of control, *XCR1.Bcl6^KO^* and *CD11c.Bcl6^KO^* mice. The plots show representative staining profiles of live, CD45⁺, lineage⁻ (CD3, CD19, CD64, B220, NK1.1), MHCII⁺, CD11c^hi^ gated single cells. Right: Absolute numbers of cDC subsets in the lungs of control, *XCR1.Bcl6^KO^*, and *CD11c.Bcl6^KO^* mice. Statistical analysis by one-way ANOVA.** P<0.01, **** P<0.0001. Data points represent values from individual mice pooled from 2-3 experiments and lines represent means +/- SD.

B. Left: Representative flow cytometry plots of blood XCR1⁺ and XCR1⁻CD11b⁺ DC subsets from control, XCR1.Bcl6^KO^ and CD11c.Bcl6^KO^. Shown are representative staining profiles of live, CD45⁺, lineage⁻ (CD3, CD19, CD64, B220, NK1.1), MHCII⁺, CD11c^hi^ gated single cells. Right: Absolute numbers of cDC1 and cDC2 in blood of control, XCR1.Bcl6^KO^, and CD11c.Bcl6^KO^ mice. Statistical analysis by one-way ANOVA, significance at * P<0.05. Data points are normalized to volume of blood drawn and represent values from individual mice pooled from 4 experiments; lines represent means +/- SD from each genotype.

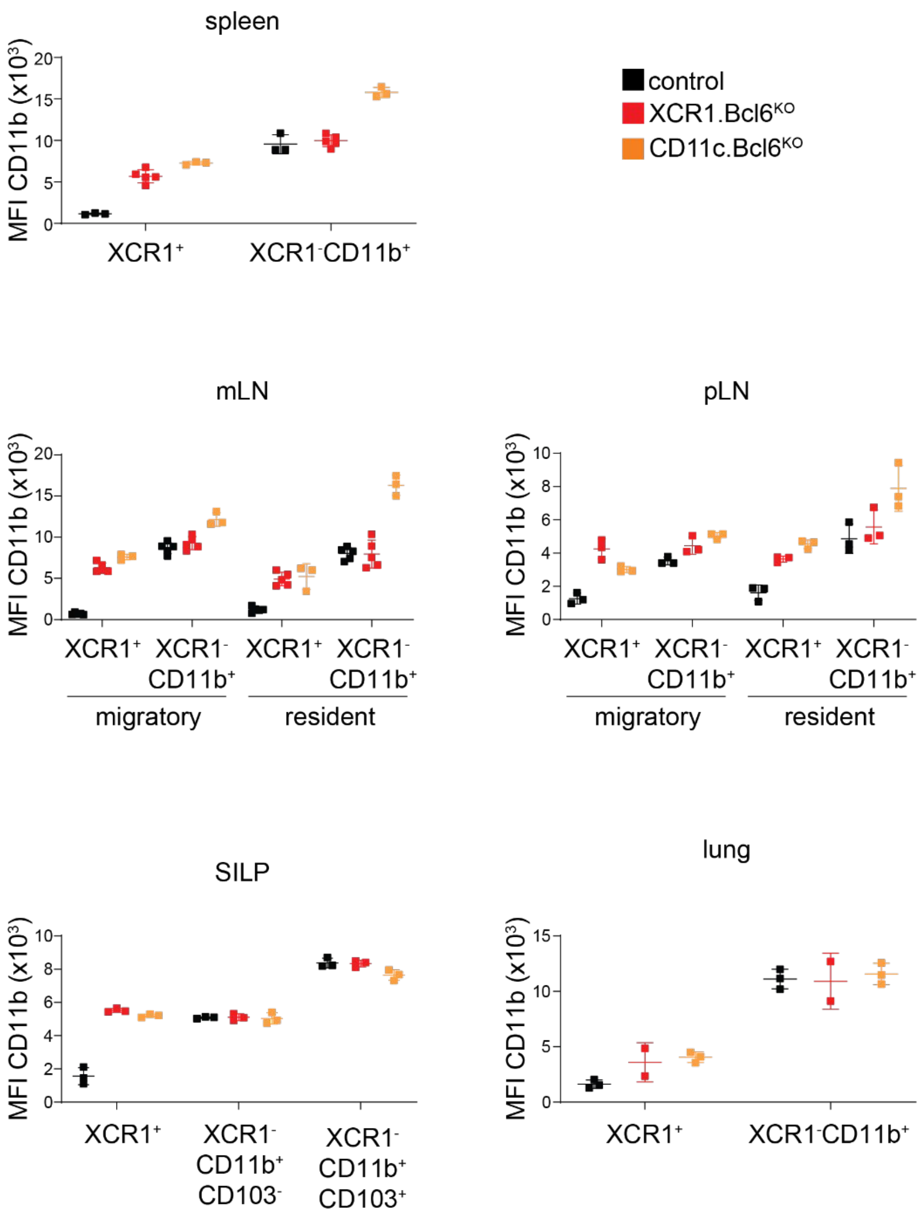


### Supplementary Figure 3

Raw values of CD11b MFI in XCR1$^+$ and XCR1$^-$CD11b$^+$ (spleen, mLN, pLN and lung) or XCR1$^+$ and XCR1$^-$CD11b$^+$CD103$^-$ and CD11b$^+$CD103$^+$ (SILP) cDC from control, *XCR1.Bcl6$^{KO}$*, and *CD11c.Bcl6$^{KO}$* mice. Note that different organs were stained using different panels and CD11b MFI values are therefore not comparable across organs. Data depicts one representative experiment of at least 3 per organ with 2-3 mice each.

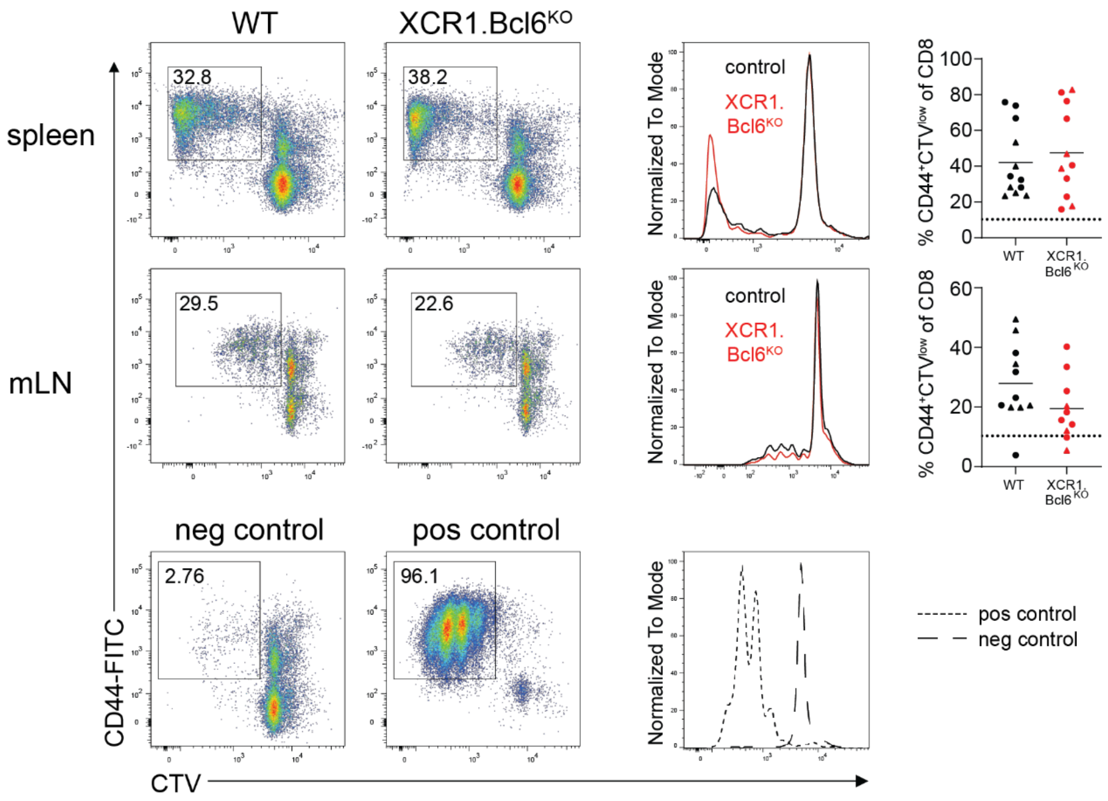


**Supplementary Figure 4**

Representative flow cytometry plots showing CTV dilution and CD44 expression on OT-I cells (gated on live, CD11c$^{-}$, CD8α$^{+}$ single cells) following 65h incubation with OVA pulsed and LPS stimulated cDC1 isolated from spleen (top) and mLN (bottom) of control and *XCR1.Bcl6$^{KO}$* mice. Data are representative of four independent experiments with 2-3 mice each.

Histograms show CTV dilution in OT-I cells after incubation with OVA pulsed cDC1 isolated from spleen and mLN of control and *XCR1.Bcl6$^{KO}$* mice. Bottom histogram shows CTV dilution in OT-I cells after incubation with anti-CD3 and anti-CD28 (positive control) and non-pulsed cDC1 (negative control).

Percentage of CD44$^{+}$ CTV$^{low}$ OT-I cells after incubation with cDC1s isolated from spleen and mLN of control and *XCR1.Bcl6$^{KO}$* mice. Data is pooled from four independent experiments. Each dot represents one mouse, solid lines represent means +/- SD and dotted lines represent the average percentage of CD44$^{+}$ CTV$^{low}$ OT-I cells from negative controls across all independent experiments.

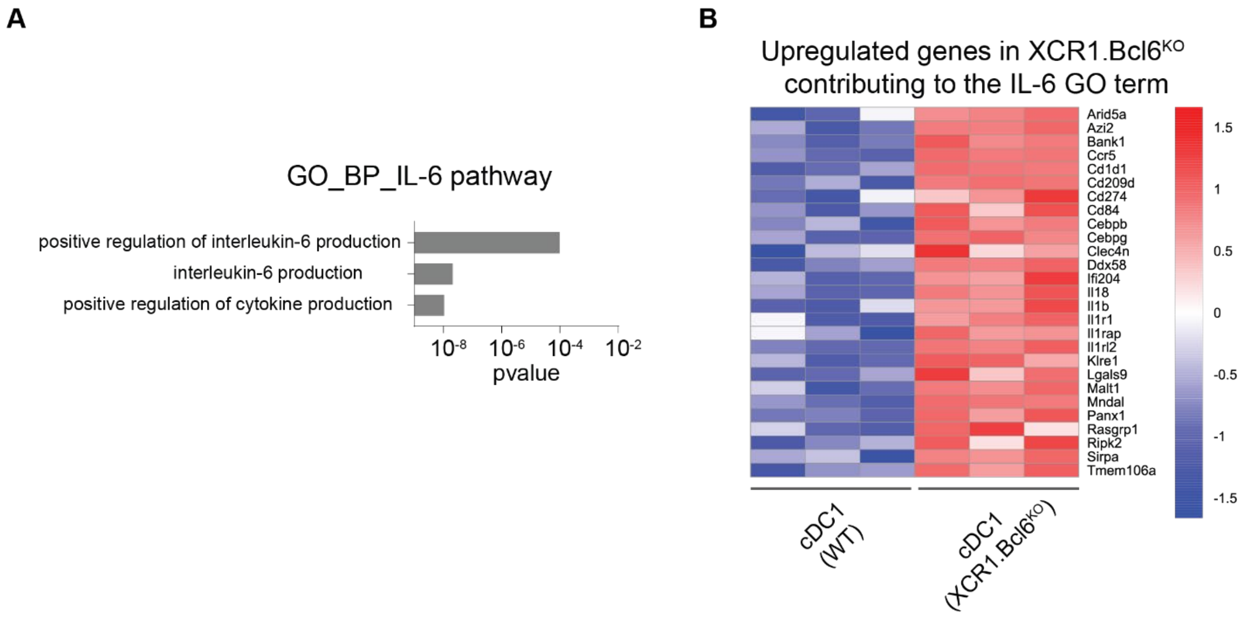


**Supplementary Figure 5**

A. IL-6 pathway gene ontology (GO) term enrichment analysis of differentially expressed genes (DEGs) in *XCR1.Bcl6*$^{KO}$ and control cDC1. The p value calculated from GO term enrichment analysis is represented on the x-axis and the y-axis shows the GO Biological Process (BP) terms. Only terms related to positive regulation of IL-6 production are depicted.
B. Heatmap of all upregulated genes in *XCR1.Bcl6*$^{KO}$ cDC1 contributing to the enriched IL-6 pathways depicted in A. The color scale represents the row Z score.

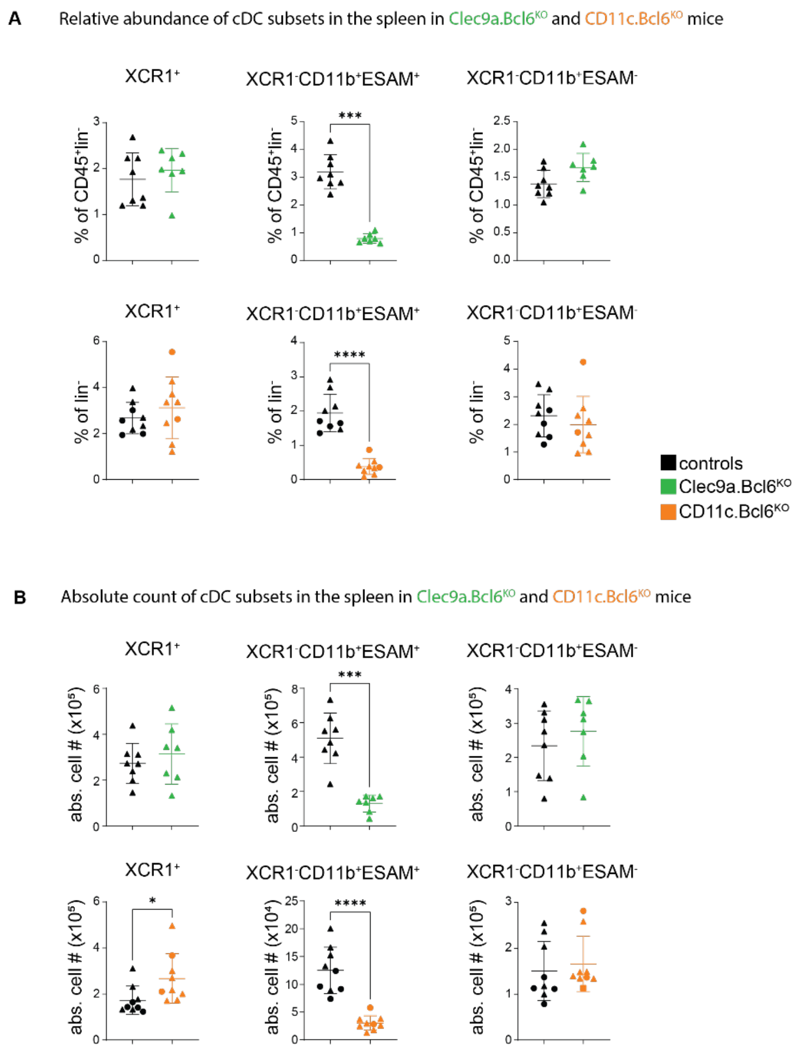


### Supplementary Figure 6

A. Relative proportions of XCR1⁺, XCR1⁻CD11b⁺ESAM$^{hi}$ and XCR1⁻CD11b⁺ESAM$^{lo}$ subsets in the spleen of Clec9a.*Bcl6$^{KO}$* (top) and CD11c.*Bcl6$^{KO}$* (bottom) mice. Cells were pre-gated on single live, CD45⁺, lineage⁻ (CD3, CD19, TER119, B220, NK1.1), MHCII⁺, CD11c$^{hi}$. Data points represent values from individual mice pooled from 2-3 experiments and lines represent means +/- SD. Statistical analysis using Mann-Whitney test, *** P<0.001 and **** P<0.0001.

B. Numbers of XCR1⁺, XCR1⁻CD11b⁺ESAM$^{hi}$ and XCR1⁻CD11b⁺ESAM$^{lo}$ subsets in the spleen of control, Clec9a.*Bcl6$^{KO}$* normalized to volume of suspension (top) and CD11c.*Bcl6$^{KO}$* shown as absolute numbers (bottom). Data points represent values from individual mice pooled from 2-3 experiments and lines represent means +/- SD. * P<0.05, *** P<0.001 and **** P<0.0001, by Mann-Whitney test.

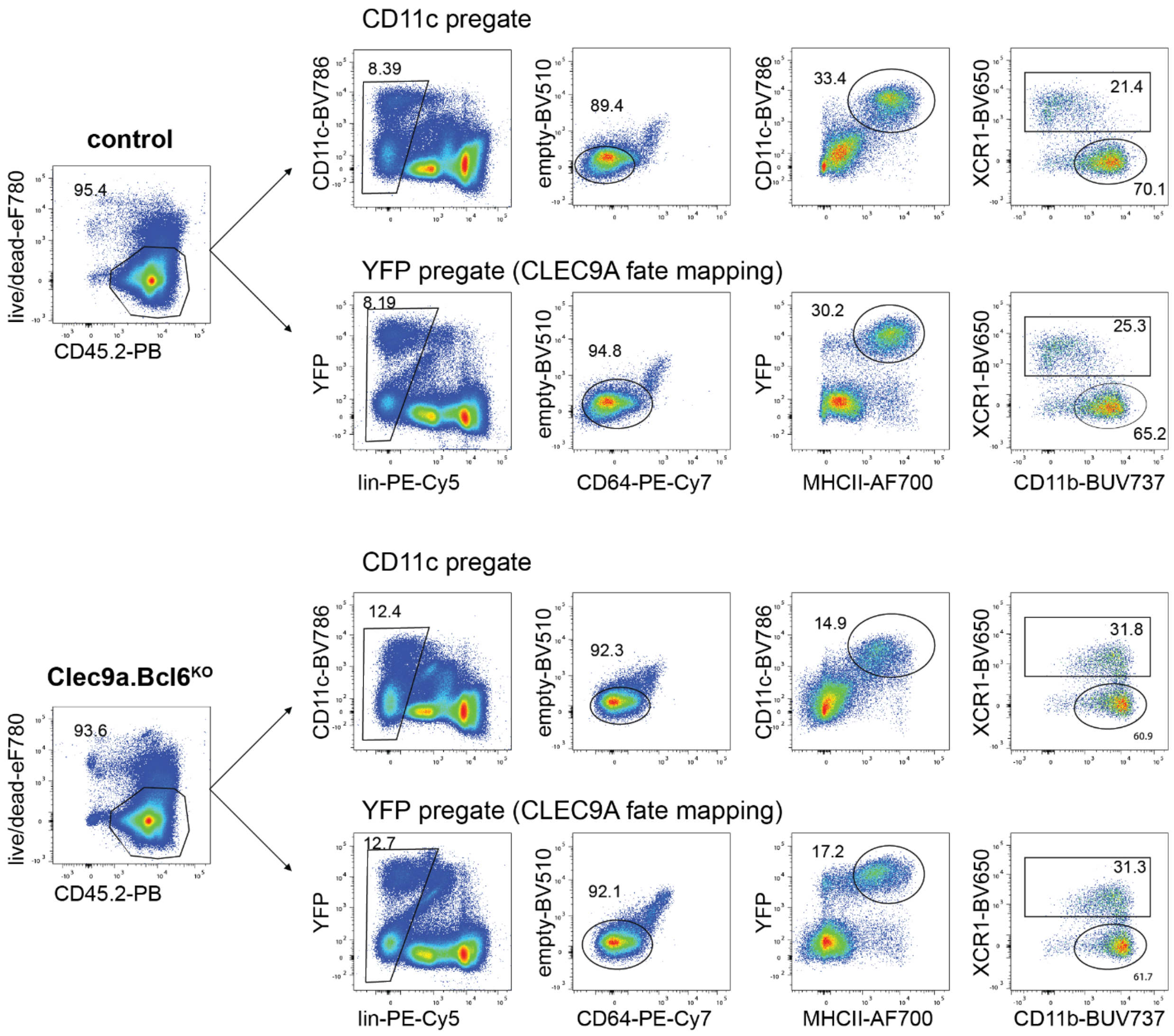


**Supplementary Figure 7**

FACS plots showing alternative gating strategies for cDCs in Clec9a.*Bcl6*$^{KO}$ and control mice. CD11c pre-gate: classical CD11c-based gating; YFP pre-gate: Gating based on *Clec9a* fate mapping (Clec9a.*Bcl6*$^{KO}$ mice contain the *R26-ROSA*$^{YFP}$ locus). Data is representative for 3 experiments with 2-3 mice each.

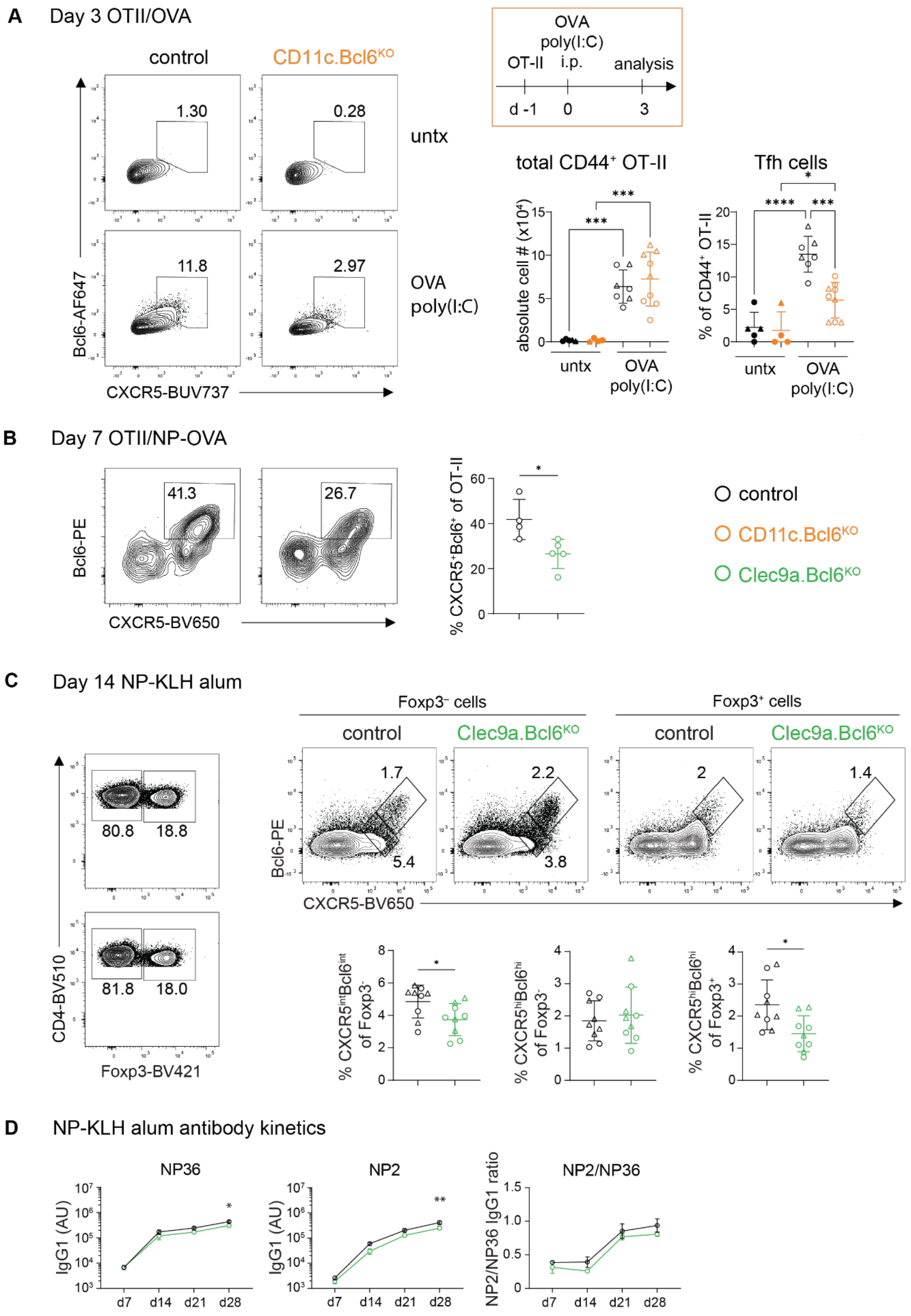

A
Day 3 OTII/OVA
control
CD11c.Bcl6KO
1.30
0.28
untx
11.8
2.97
OVA
poly(I:C)
Bcl6-AF647
CXCR5-BUV737
OVA
poly(I:C)
OT-II
i.p.
analysis
d -1
0
3
total CD44+ OT-II
absolute cell # (x10⁴)
untx
OVA
poly(I:C)
Tfh cells
% of CD44+ OT-II
B
Day 7 OTII/NP-OVA
41.3
26.7
Bcl6-PE
CXCR5-BV650
% CXCR5+Bcl6+ of OT-II
control
CD11c.Bcl6KO
Clec9a.Bcl6KO
C
Day 14 NP-KLH alum
Foxp3– cells
Foxp3+ cells
control
Clec9a.Bcl6KO
80.8
18.8
81.8
18.0
CD4-BV510
Foxp3-BV421
1.7
5.4
2.2
3.8
2
1.4
Bcl6-PE
CXCR5-BV650
% CXCR5intBcl6int of Foxp3-
% CXCR5hiBcl6hi of Foxp3-
% CXCR5hiBcl6hi of Foxp3+
D
NP-KLH alum antibody kinetics
NP36
NP2
NP2/NP36
IgG1 (AU)
NP2/NP36 IgG1 ratio
d7
d14
d21
d28

**Supplementary Figure 8**

A. Representative flow cytometric analysis of early Tfh cell induction amongst OT-II cells in the spleen of control and *CD11c.Bcl6*$^{KO}$ mice 3 days after i.p. immunization with OVA and poly(I:C) or with PBS as a control. Tfh were gated as live CD4$^+$V□5.2$^+$CD45.1$^+$CD45.2$^+$CD44$^+$Bcl6$^+$CXCR5$^+$ and the results are representative of three experiments with 3-5 mice each. Right: Numbers of total CD44$^+$ OT-II T cells (left panel), left: proportions (right panel) of OVA-specific Tfh cells in the spleens of control and *CD11c.Bcl6*$^{KO}$ mice 3 days after i.p. immunization with OVA and poly(I:C) or with PBS as a control. Statistical analysis was performed by one-way ANOVA. * P<0.05, ** P<0.01, *** P<0.001 and **** P<0.0001. Data points represent values from individual mice pooled from three experiments with 3-5 mice each in OVA/poly(I:C) immunized groups. Each experiment contained at least one untreated control per genotype; lines represent means +/- SD.

B. Bcl6-staining and Bcl6-based quantification of Tfh cells of data presented in Figure 8C, D.

C. Foxp3-staining of CD4$^+$ T cells and Bcl6-staining and Bcl6-based quantification of Tfh cells, GC Tfh cells, and Tfr cells complementary to data presented in Figure 8E to H.

D. NP-specific IgG1 antibody titers accompanying total IgG data presented in Figure 8I, J.

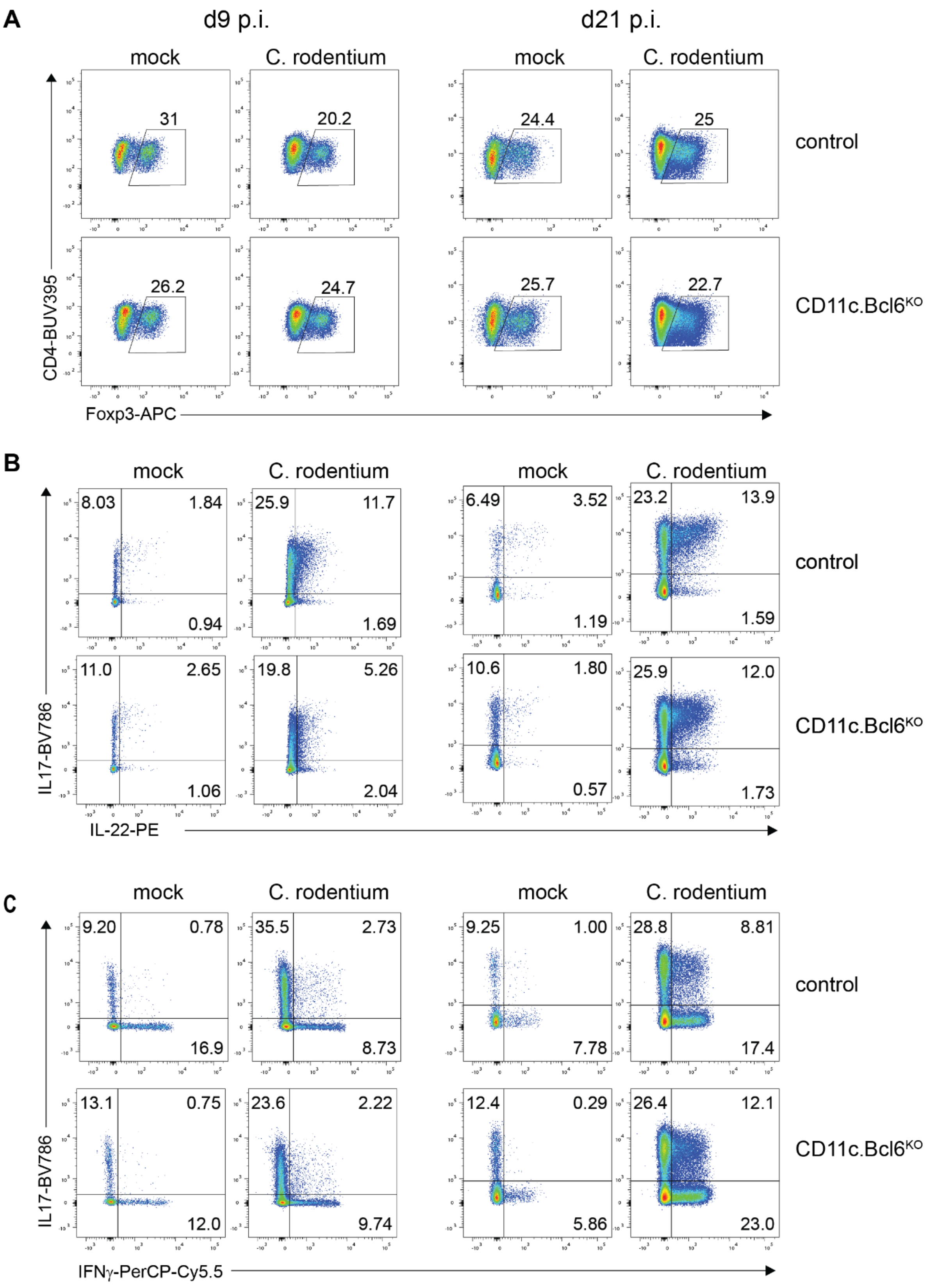
A
d9 p.i.
d21 p.i.
mock
C. rodentium
mock
C. rodentium
31
20.2
24.4
25
control
26.2
24.7
25.7
22.7
CD11c.Bcl6KO
CD4-BUV395
Foxp3-APC
B
mock
C. rodentium
mock
C. rodentium
8.03
1.84
0.94
25.9
11.7
1.69
6.49
3.52
1.19
23.2
13.9
1.59
control
11.0
2.65
1.06
19.8
5.26
2.04
10.6
1.80
0.57
25.9
12.0
1.73
CD11c.Bcl6KO
IL17-BV786
IL-22-PE
C
mock
C. rodentium
mock
C. rodentium
9.20
0.78
16.9
35.5
2.73
8.73
9.25
1.00
7.78
28.8
8.81
17.4
control
13.1
0.75
12.0
23.6
2.22
9.74
12.4
0.29
5.86
26.4
12.1
23.0
CD11c.Bcl6KO
IL17-BV786
IFNγ-PerCP-Cy5.5

**Supplementary Figure 9**

A. Representative flow cytometry plots showing Foxp3$^{+}$ CD4$^{+}$ T cells (gated on live, CD45$^{+}$, CD3$^{+}$TCRb$^{+}$ singlets) in the cLP of mock and C. rodentium infected control and CD11c.Bcl6$^{KO}$ mice at day 9 (left) or day 21 (right) post inoculation (used for main article Figure 10).

B. Representative flow cytometry analysis showing IL-17A$^{+}$IL-22$^{+}$ CD4$^{+}$ T cells (gated on live, CD45$^{+}$, CD3$^{+}$TCRb$^{+}$ singlets) in the cLP of mock and C. rodentium infected control and CD11c.Bcl6$^{KO}$ mice at day 9 (left) or day 21 (right) post inoculation (used for main article Figure 10).

C. Representative flow cytometry plots showing IL-17A$^{+}$IFNg$^{+}$ CD4$^{+}$ T cells (gated on live, CD45+, CD3$^{+}$TCRb$^{+}$ singlets) in the cLP of mock and C. rodentium infected control and CD11c.Bcl6$^{KO}$ mice at day 9 (left) or day 21 (right) post inoculation (used for main article Figure 10).

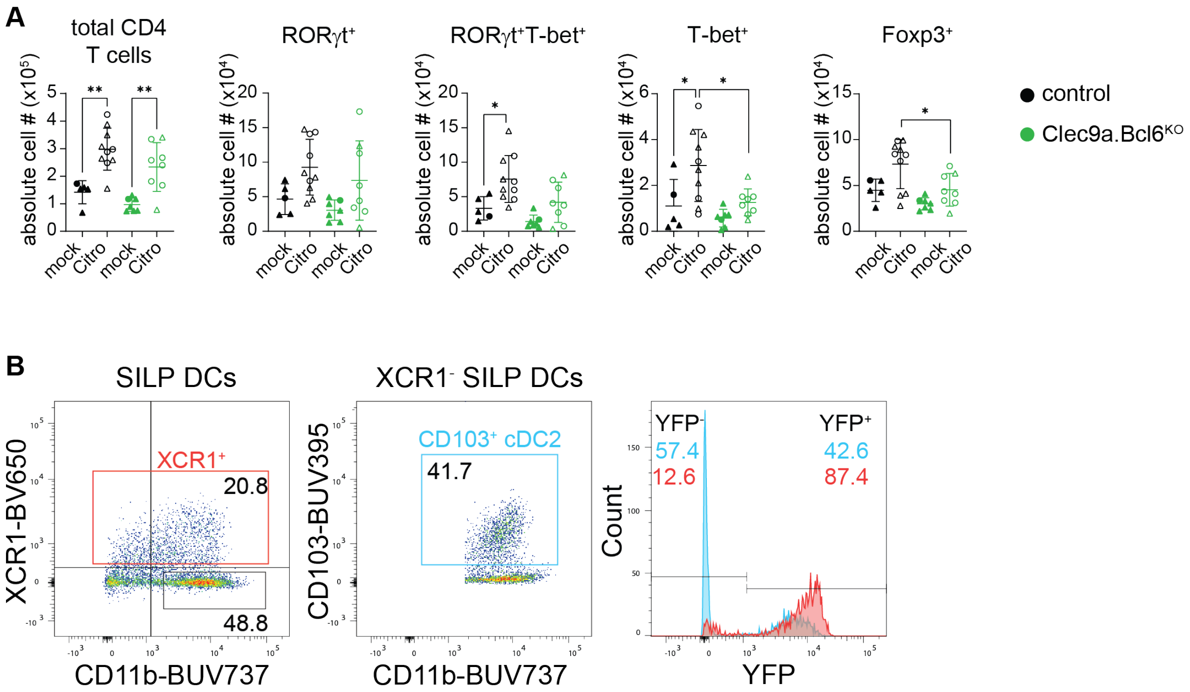


**Supplementary Figure 10**

A. Numbers of total CD4⁺ T cells, RORgt⁺, RORgt⁺T-bet⁺, T-bet⁺, and Foxp3⁺ CD4⁺ T cells in the cLP of mock and *C. rodentium* infected control and *Clec9a.Bcl6*$^{KO}$ mice at day 9 post inoculation. Statistical analysis performed by one-way ANOVA, * P<0.05, ** P<0.01, *** P<0.001 and **** P<0.0001. Data representative of 3 independent experiments with 1-3 mice each.

B. Left: Flow cytometry plot showing XCR1⁺ and XCR1⁻CD11b⁺ gating on SILP DCs, pre-gated on live, lineage⁻ (CD3, CD19, CD64, B220, NK1.1), CD11c⁺, MHCII⁺ single cells; right: Sub-gate on XCR1⁻CD11b⁺ cells. Histogram shows YFP-expression in XCR1+ cDC1 (red) and XCR1⁻CD11b⁺CD103⁺ cDC2 (blue). Data representative of 2 experiments with 3 mice each.
