## Supplementary material for "Genomic deletion of Bcl6 differentially affects conventional dendritic cell subsets and compromises Tfh/Tfr/Th17 cell responses": Suppl Table 4

| Product | Company | Cat. No. | Dilution | Product link |
| --- | --- | --- | --- | --- |
| Collagenase, Type IV, C. histolyticum | US biological | C7511-70 | 0.5 mg/ml | https://www.usbio.net/molecular-biology/C7511-70/collagenase-type-iv-c-histolyticum |
| DNase I grade II | Roche | 10104159001 | 30 µg/ml | https://www.sigmaaldrich.com/DK/en/product/roche/10104159001 |
| HBSS (10X), no calcium, no magnesium, phenol red | Gibco/ThermoFisher | 14180046 | | https://www.thermofisher.com/order/catalog/product/14180046?tsid=Email_POE_OC_OrderConfirm%20%20_SKULINK |
| RPMI 1640 Medium | Gibco/ThermoFisher | 11875093 | | https://www.thermofisher.com/order/catalog/product/11875093?SID=srch-srp-11875093 |
| DPBS, calcium, magnesium | Gibco/ThermoFisher | 14040174 | | https://www.thermofisher.com/order/catalog/product/14040174?SID=srch-hj-14040174 |
| DPBS, no calcium, no magnesium | Gibco/ThermoFisher | 14190250 | | https://www.thermofisher.com/order/catalog/product/14190250?SID=srch-srp-14190250 |
| Penicillin-Streptomycin-Glutamin 100x | Gibco/ThermoFisher | 10378016 | | https://www.thermofisher.com/order/catalog/product/10378016?SID=srch-srp-10378016 |
| EDTA (0.5 M), pH 8.0, RNase-free | Gibco/ThermoFisher | AM9260G | | https://www.thermofisher.com/order/catalog/product/AM9260G?SID=srch-srp-AM9260G |
| 2-Mercaptoethanol (50 mM) | Gibco/ThermoFisher | 31350010 | | https://www.thermofisher.com/order/catalog/product/31350010?SID=srch-srp-31350010 |
| Viralex™ Foetal Calf Serum | PAA Laboratories GmbH | A15-651 | | |
| Triton X-100 | Sigma | X-100 | | https://www.sigmaaldrich.com/CA/en/product/sial/x100 |
| PFA | Thermo Fisher | J19943.K2 | | https://www.thermofisher.com/order/catalog/product/J19943.K2?SID=srch-hj-J19943.K2 |
| HEPES (1M) | Gibco/ThermoFisher | 15630056 | | https://www.thermofisher.com/order/catalog/product/15630056?SID=srch-srp-15630056 |
| Liberase™ TM Research Grade | Roche | LIBTM-RO | 58 ug/ml | https://www.sigmaaldrich.com/DK/en/product/roche/libtmro?utm_source=google&utm_medium=cpc&utm_campaign=8809292844&utm_content=88946462815&gclid=CjwKCAiAlcyuBhBnEiwAOGZ2SxdeMCvw56qgz2Q-N1PfcUXLuywrJ2YOXNk9vWObKdOzPSV1h9dRTRoC-HQQAvD_BwE |
| Percoll® | Cytiva/GE Healthcare/SigmaA | GE17-0891-01 | | https://www.sigmaaldrich.com/DK/en/product/sigma/ge17089101 |
| Dynabeads™ Mouse DC (Dendritic Cell) Enrichment | Invitrogen/ThermoFisher | 11429D | | https://www.thermofisher.com/order/catalog/product/11429D |
| Buffer RLT | QIAGEN | 79216 | | https://www.qiagen.com/us/products/discovery-and-translational-research/lab-essentials/buffers-reagents/buffer-rlt |
| QIAGEN RNeasy Mini kit | QIAGEN | 74106 | | https://www.qiagen.com/us/products/discovery-and-translational-research/dna-rna-purification/rna-purification/total-rna/rneasy-kits?catno=74106 |
| RNase-Free DNase Set | QIAGEN | 79254 | | https://www.qiagen.com/us/products/discovery-and-translational-research/lab-essentials/enzymes/rnase-free-dnase-set |
| Ovation® Ultralow System V2 | Tecan | 0344NB | | https://shop.tecan.com/va/en/Reagents-&-Kits/Next-Gen-Sequencing/Ovation%C2%AE-Ultralow-System-V2,-32-reactions/p/0344NB-32 |
| Ovation® RNA-Seq System V2 | Tecan | 7102 | | https://shop.tecan.com/va/en/product/7102-32/OvationRNA-Seq-System-V2-32-reactions |
| AMPure XP Bead-Based Reagent | Beckman Coulter | A63880 | | https://www.beckman.com/reagents/genomic/cleanup-and-size-selection/pcr/a63880 |
| Dynabeads™ Untouched™ Mouse T Cells Kit | Invitrogen/ThermoFisher | 11413D | | https://www.thermofisher.com/order/catalog/product/11413D |
| Dynabeads™ Untouched™ Mouse CD8 Cells Kit | Invitrogen/ThermoFisher | 11417D | | https://www.thermofisher.com/order/catalog/product/11417D |
| Dynabeads™ Untouched™ Mouse CD4 Cells Kit | Invitrogen/ThermoFisher | 11415D | | https://www.thermofisher.com/order/catalog/product/11417D |
| CellTrace™ Violet Cell Proliferation Kit, for flow cytometry | Invitrogen/ThermoFisher | C34571 | | https://www.thermofisher.com/order/catalog/product/C34571?SID=srch-hj-C34571 |
| Fingolimod (hydrochloride) | Cayman Chemical | 10006292 | 20 µg/mou | https://www.caymanchem.com/product/10006292/fingolimod-(hydrochloride) |
| OVA EndoFit™ | InvivoGen | vac-pova | | https://www.invivogen.com/endofit-ovalbumin |
| LPS-RS | InvivoGen | tlrl-rslps | | https://www.invivogen.com/lps-rs |
| Albumin from chicken egg white | SigmaAldrich | A5503 | | https://www.sigmaaldrich.com/DK/en/product/sigma/a5503 |
| Polyinosinic–polycytidylic acid sodium salt | SigmaAldrich | P0913 | 100 ug/mouse | |
| $NP_{17}$-OVA ((#N-5051-10-BS Biosearch Technologies/BioCat) | | | 50 µg | |
| Imject Alum (Thermo Fisher)). | | | | |
| 50µg $NP_{32}$-KLH (#N-5060-5, Biosearch Technologies/BioCat) | | | | |
| CD16/CD32 Monoclonal Antibody (93), eBioscience™ | Invitrogen/ThermoFisher | 14-0161-86 | 1:100 | https://www.thermofisher.com/antibody/product/CD16-CD32-Antibody-clone-93-Monoclonal/14-0161-86 |
| LIVE/DEAD™ Fixable Near-IR Dead Cell Stain Kit | Invitrogen/ThermoFisher | L34976 | 1:1000 | https://www.thermofisher.com/order/catalog/product/L34976?SID=srch-srp-L34976 |
| LIVE/DEAD™ Fixable Near IR (780) Viability Kit | Invitrogen/ThermoFisher | L34994 | 1:1000 | https://www.thermofisher.com/order/catalog/product/L34994?SID=srch-srp-L34994 |
| DRAQ7 | Nordic Biosite/BioLegend | 424001 | Dirk | https://www.biolegend.com/fr-ch/products/draq7-9628?GroupID=GROUP22 |
| Recombinant Mouse IL-23 Protein | R&D systems | 1887-ML-010/ | 40 ng/ml | |
| PMA | Sigma-Aldrich | P1585-1MG | 50 ng/ml | https://www.sigmaaldrich.com/DK/en/product/sigma/p1585 |
| Ionomycin calcium salt from Streptomyces conglobatus | Sigma-Aldrich | I0634-1MG | 750 ng/ml | https://www.sigmaaldrich.com/DK/en/product/sigma/i0634 |
| BD GolgiStop™ Protein Transport Inhibitor (Containing Monensin) | BD Biosciences | 554724 | 1:10000 | https://www.bdbiosciences.com/en-dk/products/reagents/flow-cytometry-reagents/research-reagents/single-color-antibodies-ruo/protein-transport-inhibitor-containing-monensin.554724 |
| eBioscience™ Foxp3 / Transcription Factor Staining Buffer Set | Invitrogen/ThermoFisher | 00-5523-00 | N/A | https://www.thermofisher.com/order/catalog/product/00-5523-00 |
| BD OptEIA™ Mouse IL-6 ELISA Set | BD Biosciences | 555240 | N/A | https://www.bdbiosciences.com/en-ca/products/reagents/immunoassay-reagents/elisa/elisa-kits/mouse-il-6-elisa-set.555240 |
| **Antibodies** | | | | |
| **In vitro** | | | | |
| CD3 Monoclonal Antibody (17A2), Functional Grade, eBioscience™ | Invitrogen/ThermoFisher | 16-0032-82 | 10 ug/ml | https://www.thermofisher.com/antibody/product/CD3-Antibody-clone-17A2-Monoclonal/16-0032-82 |
| CD28 Monoclonal Antibody (37.51), eBioscience™ | Invitrogen/ThermoFisher | 14-0281-82 | 1 ug/ml | https://www.thermofisher.com/antibody/product/CD28-Antibody-clone-37-51-Monoclonal/14-0281-82 |
| **Imaging** | | | | |
| BV421 anti-mouse B220/CD45R (clone RA3-6B2) | BD Biosciences | 562922 | 0.2mg/ml | https://www.bdbiosciences.com/en-us/products/reagents/flow-cytometry-reagents/research-reagents/single-color-antibodies-ruo/bv421-rat-anti-mouse-cd45r-b220.562922 |
| APC anti-mouse CD283 (TLR3), clone 11F8 | Nordic Biosite/BioLegend | 141904 | 0.2mg/ml | https://nordicbiosite.com/product/141904/PE-antimouse-CD283-TLR3 |
| Alexa Fluor® 488 anti-mouse CD11c (clone N418) | Nordic Biosite/BioLegend | 117311 | 0.2mg/ml | https://www.biolegend.com/de-at/products/alexa-fluor-488-anti-mouse-cd11c-antibody-2702 |
| PE/Dazzle™ 594 anti-mouse CD172a (SIRPα), clone P84 | Nordic Biosite/BioLegend | 144016 | 0.2mg/ml | https://www.biolegend.com/en-us/products/pe-dazzle-594-anti-mouse-cd172a-sirpalpha-antibody-10804 |
| unlabeled anti-GFP (polyclonal) | Invitrogen | A10262 | 2mg/ml | https://www.thermofisher.com/antibody/product/GFP-Antibody-Polyclonal/A10262 |
| Goat anti-Chicken IgY (H+L) Secondary Antibody, Alexa Fluor™ 488, polyclonal | Jackson Immunoresearch | 703-546-155 | 1.5mg/ml | https://www.jacksonimmuno.com/catalog/products/703-546-155 |
| **Flow Cytometry** | | | | |
| NK1.1 Monoclonal Antibody (PK136), PE-Cyanine7, eBioscience™ | Invitrogen/ThermoFisher | 25-5941-82 | 1:150 | https://www.thermofisher.com/antibody/product/NK1-1-Antibody-clone-PK136-Monoclonal/25-5941-82 |
| CD3e Monoclonal Antibody (145-2C11), PE-Cyanine7, eBioscience™ | Invitrogen/ThermoFisher | 25-0031-82 | 1:200 | https://www.thermofisher.com/antibody/product/CD3e-Antibody-clone-145-2C11-Monoclonal/25-0031-82 |
| CD45R (B220) Monoclonal Antibody (RA3-6B2), PE-Cyanine7, eBioscience™ | Invitrogen/ThermoFisher | 25-0452-82 | 1:200 | https://www.thermofisher.com/antibody/product/CD45R-B220-Antibody-clone-RA3-6B2-Monoclonal/25-0452-82 |
| CD19 Monoclonal Antibody (eBio1D3 (1D3)), PE-Cyanine7, eBioscience™ | Invitrogen/ThermoFisher | 25-0193-82 | 1:100 | https://www.thermofisher.com/antibody/product/CD19-Antibody-clone-eBio1D3-1D3-Monoclonal/25-0193-82 |
| PE/Cyanine7 anti-mouse CD64 (FcγRI), clone X54-5/7.1 | Nordic Biosite/BioLegend | 139314 | 1:100 | https://nordicbiosite.com/product/139314/PECyanine7-antimouse-CD64-FcgRI |
| NK1.1 APC-Cy7 | | | | |
| BD Horizon™ BUV737 Rat Anti-Mouse CD3 Molecular Complex, clone 17A2 | BD Biosciences | 612803 | 1:200 | https://www.bdbiosciences.com/en-dk/products/reagents/flow-cytometry-reagents/research-reagents/single-color-antibodies-ruo/buv737-rat-anti-mouse-cd3-molecular-complex.612803 |
| CD3 APC-Cy7 | | | | |
| B220 APC-Cy7 | | | | |
| CD19 APC-Cy7 | | | | |
| CD19 BUV737 | | | | |
| APC-Cy7 anti-mouse TER119 | | | | |
| APC anti-mouse CD11c, clone N418 | Nordic Biosite/BioLegend | 117310 | 1:100 | https://nordicbiosite.com/product/117310/APC-antimouse-CD11c |
| BD Horizon™ BV711 Hamster Anti-Mouse CD11c, clone HL3 | BD Biosciences | 563048 | 1:200 | https://www.bdbiosciences.com/en-dk/products/reagents/flow-cytometry-reagents/research-reagents/single-color-antibodies-ruo/bv711-hamster-anti-mouse-cd11c.563048 |
| PerCP-Cy5.5 anti-mouse CD11c | | | | |
| FITC anti-mouse I-A/I-E, clone M5/114.15.2 | Nordic Biosite/BioLegend | 107606 | 1:200 | https://nordicbiosite.com/product/107606/FITC-antimouse-IAIE |
| MHC-II BUV661 | | | | |
| MHC Class II (I-A/I-E) Monoclonal Antibody (M5/114.15.2), Alexa Fluor™ 700, eBioscience™ | Invitrogen/ThermoFisher | 56-5321-82 | 1:200 | https://www.thermofisher.com/antibody/product/MHC-Class-II-I-A-I-E-Antibody-clone-M5-114-15-2-Monoclonal/56-5321-82?gclid=CjwKCAiA8sauBhB3EiwAruTRJuPUcPzyotr4ezWZHeSPxEWWRmS_Fv0EB4H3_l9xkOynZalxZsr-iBoCzAoQAvD_BwE&ef_id=CjwKCAiA8sauBhB3EiwAruTRJuPUcPzyotr4ezWZHeSPxEWWRmS_Fv0EB4H3_l9xkOynZalxZsr-iBoCzAoQAvD_BwE:G:s&s_kwcid=AL!3652!3!600168785949!!!g!!!1454324556!63404918784&cid=bid_pca_frg_r01_co_cp1359_pjt0000_bid00000_0se_gaw_dy_pur_con&gad_source=1 |
| CD11b Monoclonal Antibody (M1/70), APC-Cyanine7 | Invitrogen/ThermoFisher | A15390 | 1:400 | https://www.thermofisher.com/antibody/product/CD11b-Antibody-clone-M1-70-Monoclonal/A15390 |
| BD Horizon™ BUV737 Rat Anti-CD11b, clone M1/70 | BD Biosciences | 564443 | 1:400 | https://www.bdbiosciences.com/en-dk/products/reagents/flow-cytometry-reagents/research-reagents/single-color-antibodies-ruo/buv737-rat-anti-cd11b.612801 |
| FITC CD11b | | | | |
| APC-R700 CD11b | | | | |
| PE anti-mouse/rat XCR1, clone ZET | Nordic Biosite/BioLegend | 148204 | 1:100 | https://nordicbiosite.com/product/148204/PE-antimouserat-XCR1 |
| Brilliant Violet 421™ anti-mouse/rat XCR1, clone ZET | Nordic Biosite/BioLegend | 148216 | 1:200 | https://nordicbiosite.com/product/148216/Brilliant-Violet-421tm-antimouserat-XCR1 |
| BD Horizon™ BV421 Rat Anti-Mouse CD103, clone 290 | BD Biosciences | 562771 | 1:100 | https://www.bdbiosciences.com/en-dk/products/reagents/flow-cytometry-reagents/research-reagents/single-color-antibodies-ruo/bv421-rat-anti-mouse-cd103.562771 |
| BD Horizon™ BV786 Rat Anti-Mouse CD103, clone M290 | BD Biosciences | 564322 | 1:200 | https://www.bdbiosciences.com/en-dk/products/reagents/flow-cytometry-reagents/research-reagents/single-color-antibodies-ruo/bv786-rat-anti-mouse-cd103.564322 |
| CD24 Monoclonal Antibody (M1/69), PE-Cyanine7, eBioscience™ | Invitrogen/ThermoFisher | 25-0242-82 | 1:200 | https://www.thermofisher.com/antibody/product/CD24-Antibody-clone-M1-69-Monoclonal/25-0242-82?gclid=CjwKCAiA8sauBhB3EiwAruTRJozLF8pKH6b2TTqcMGJde35U1uK5UmhwWkqD0lIKMIjYObisJQZ9SxoCn-0QAvD_BwE&ef_id=CjwKCAiA8sauBhB3EiwAruTRJozLF8pKH6b2TTqcMGJde35U1uK5UmhwWkqD0lIKMIjYObisJQZ9SxoCn-0QAvD_BwE:G:s&s_kwcid=AL!3652!3!600168785949!!!g!!!1454324556!63404918784&cid=bid_pca_frg_r01_co_cp1359_pjt0000_bid00000_0se_gaw_dy_pur_con&gad_source=1 |
| BD Horizon™ BV605 Rat Anti-Mouse CD8a, clone 53-6.7 | BD Biosciences | 563152 | 1:100 | https://www.bdbiosciences.com/en-dk/products/reagents/flow-cytometry-reagents/research-reagents/single-color-antibodies-ruo/bv605-rat-anti-mouse-cd8a.563152 |
| APC anti-mouse CD283 (TLR3), clone 11F8 | Nordic Biosite/BioLegend | 141906 | 1:100 | https://nordicbiosite.com/product/141906/APC-antimouse-CD283-TLR3 |
| PE anti-mouse CD283 (TLR3), clone 11F8 | Nordic Biosite/BioLegend | 141904 | 1:100 | https://nordicbiosite.com/product/141904/PE-antimouse-CD283-TLR3 |
| BD Horizon™ BUV395 Rat Anti-Mouse CD45, clone 30-F11 | BD Biosciences | 564279 | 1:500 | https://www.bdbiosciences.com/en-dk/products/reagents/flow-cytometry-reagents/research-reagents/single-color-antibodies-ruo/buv395-rat-anti-mouse-cd45.564279 |
| BD Horizon™ V500 Rat Anti-Mouse CD45, clone 30-F11 | BD Biosciences | 561487 | 1:200 | https://www.bdbiosciences.com/en-dk/products/reagents/flow-cytometry-reagents/research-reagents/single-color-antibodies-ruo/v500-rat-anti-mouse-cd45.561487 |
| PE/Dazzle™ 594 anti-mouse CD172a (SIRPα), clone P84 | Nordic Biosite/BioLegend | 144016 | 1:100 | https://nordicbiosite.com/product/144016/PEDazzletm-594-antimouse-CD172a-SIRPa |
| BD Pharmingen™ Alexa Fluor® 647 Rat Anti-Mouse ESAM, clone 1G8 | BD Biosciences | 567162 | 1:100 | https://www.bdbiosciences.com/en-dk/products/reagents/flow-cytometry-reagents/research-reagents/single-color-antibodies-ruo/alexa-fluor-647-rat-anti-mouse-esam.567162 |
| PE anti-mouse ESAM, clone 1G8 | Nordic Biosite/BioLegend | 136203 | 1:100 | https://nordicbiosite.com/product/136203/PE-antimouse-ESAM |
| PE-Cy7 anti-mouse ESAM | | | | |
| PE anti-IRF4, clone IRF4.3E4 | Nordic Biosite/BioLegend | 646404 | 1:200 | https://nordicbiosite.com/product/646404/PE-antiIRF4 |
| IRF4 Monoclonal Antibody (3E4), eFluor™ 660, eBioscience™ | Invitrogen/ThermoFisher | 50-9858-82 | 1:100 | https://www.thermofisher.com/antibody/product/IRF4-Antibody-clone-3E4-Monoclonal/50-9858-82 |
| IRF8 Monoclonal Antibody (V3GYWCH), PerCP-eFluor™ 710, eBioscience™ | Invitrogen/ThermoFisher | 46-9852-82 | 1:300 | https://www.thermofisher.com/antibody/product/IRF8-Antibody-clone-V3GYWCH-Monoclonal/46-9852-82 |
| PE anti-mouse CD370 (CLEC9A, DNGR1), clone /H11 | Nordic Biosite/BioLegend | 143504 | 1:200 | https://nordicbiosite.com/product/143504/PE-antimouse-CD370-CLEC9A-DNGR1 |
| BD Pharmingen™ FITC Mouse Anti-Mouse Vβ 5.1, 5.2 T-Cell Receptor, clone MR9-4 | BD Biosciences | 553189 | 1:200 | https://www.bdbiosciences.com/en-dk/products/reagents/flow-cytometry-reagents/research-reagents/single-color-antibodies-ruo/fitc-mouse-anti-mouse-v-5-1-5-2-t-cell-receptor.553189 |
| PerCp-Cy5.5 CD44 | | | | |
| BD Pharmingen™ FITC Rat Anti-Mouse CD44, clone IM7 | BD Biosciences | 561859 | 1:200 | https://www.bdbiosciences.com/en-dk/products/reagents/flow-cytometry-reagents/research-reagents/single-color-antibodies-ruo/fitc-rat-anti-mouse-cd44.561859 |
| APC CD44 | | | | |
| BD Horizon™ BUV737 Hamster Anti-Mouse CD69, H1.2F3 | BD Biosciences | 564684 | 1:200 | https://www.bdbiosciences.com/en-dk/products/reagents/flow-cytometry-reagents/research-reagents/single-color-antibodies-ruo/buv737-hamster-anti-mouse-cd69.612793 |
| PE CD45.2 | | | | |
| BV421 CD45.2 | | | | |
| BD Pharmingen™ Alexa Fluor® 647 Mouse anti-Bcl-6, clone K112-91 | BD Biosciences | 561525 | 1:50 | https://www.bdbiosciences.com/en-dk/products/reagents/flow-cytometry-reagents/research-reagents/single-color-antibodies-ruo/alexa-fluor-647-mouse-anti-bcl-6.561525 |
| PE Bcl6 | | | | |
| BD Pharmingen™ APC-Cy™7 Mouse Anti-Mouse CD45.1, A20 | BD Biosciences | 560579 | 1:100 | https://www.bdbiosciences.com/en-dk/products/reagents/flow-cytometry-reagents/research-reagents/single-color-antibodies-ruo/apc-cy-7-mouse-anti-mouse-cd45-1.560579 |
| BD Horizon™ BUV395 Rat Anti-Mouse CD4, clone GK1.5 | BD Biosciences | 563790 | 1:200 | https://www.bdbiosciences.com/en-dk/products/reagents/flow-cytometry-reagents/research-reagents/single-color-antibodies-ruo/buv395-rat-anti-mouse-cd4.563790 |
| BD Horizon™ BV480 Rat Anti-Mouse CD4, clone RM4-5 | BD Biosciences | 565634 | 1:200 | https://www.bdbiosciences.com/en-dk/products/reagents/flow-cytometry-reagents/research-reagents/single-color-antibodies-ruo/bv480-rat-anti-mouse-cd4.565634 |
| BD Horizon™ BV786 Rat Anti-Mouse IL-17A, clone TC11-18H10 | BD Biosciences | 564171 | 1:200 | https://www.bdbiosciences.com/en-dk/products/reagents/flow-cytometry-reagents/research-reagents/single-color-antibodies-ruo/bv786-rat-anti-mouse-il-17a.564171 |
| CD90.2 (Thy-1.2) Monoclonal Antibody (53-2.1), FITC, eBioscience™ | Invitrogen/ThermoFisher | 11-0902-82 | 1:200 | https://www.thermofisher.com/antibody/product/CD90-2-Thy-1-2-Antibody-clone-53-2-1-Monoclonal/11-0902-82 |
| IFN gamma Monoclonal Antibody (XMG1.2), PerCP-Cyanine5.5, eBioscience™ | Invitrogen/ThermoFisher | 45-7311-82 | 1:200 | https://www.thermofisher.com/antibody/product/IFN-gamma-Antibody-clone-XMG1-2-Monoclonal/45-7311-82?gclid=CjwKCAiA8sauBhB3EiwAruTRJnniycZJtAZpqcf6AK_3mqQhMV_CGshb99ybQ_XjW_cM1HaEj9itbBoCQAYQAvD_BwE&ef_id=CjwKCAiA8sauBhB3EiwAruTRJnniycZJtAZpqcf6AK_3mqQhMV_CGshb99ybQ_XjW_cM1HaEj9itbBoCQAYQAvD_BwE:G:s&s_kwcid=AL!3652!3!600168785949!!!g!!!1454324556!63404918784&cid=bid_pca_frg_r01_co_cp1359_pjt0000_bid00000_0se_gaw_dy_pur_con&gad_source=1 |
| ROR gamma (t) Monoclonal Antibody (B2D), PE-eFluor™ 610, eBioscience™ | Invitrogen/ThermoFisher | 61-6981-82 | 1:200 | https://www.thermofisher.com/antibody/product/ROR-gamma-t-Antibody-clone-B2D-Monoclonal/61-6981-82 |
| IL-22 Monoclonal Antibody (1H8PWSR), PE, eBioscience™ | Invitrogen/ThermoFisher | 12-7221-82 | 1:200 | https://www.thermofisher.com/antibody/product/IL-22-Antibody-clone-1H8PWSR-Monoclonal/12-7221-82?gclid=CjwKCAiA8sauBhB3EiwAruTRJn_4oP1jbKgAYIbCGAS6cth24oxGLdFMBrCI0iqGBsnVPrzmK1ig9RoCZwEQAvD_BwE&ef_id=CjwKCAiA8sauBhB3EiwAruTRJn_4oP1jbKgAYIbCGAS6cth24oxGLdFMBrCI0iqGBsnVPrzmK1ig9RoCZwEQAvD_BwE:G:s&s_kwcid=AL!3652!3!600168785949!!!g!!!1454324556!63404918784&cid=bid_pca_frg_r01_co_cp1359_pjt0000_bid00000_0se_gaw_dy_pur_con&gad_source=1 |
| T-bet Monoclonal Antibody (eBio4B10 (4B10)), PE-Cyanine7, eBioscience™ | Invitrogen/ThermoFisher | 25-5825-82 | 1:200 | https://www.thermofisher.com/antibody/product/T-bet-Antibody-clone-eBio4B10-4B10-Monoclonal/25-5825-82 |
| FOXP3 Monoclonal Antibody (FJK-16s), APC, eBioscience™ | Invitrogen/ThermoFisher | 17-5773-82 | 1:200 | https://www.thermofisher.com/antibody/product/FOXP3-Antibody-clone-FJK-16s-Monoclonal/17-5773-82 |
| FOXP3 Monoclonal Antibody (FJK-16s), Brilliant Violet™ 421, eBioscience™ | Invitrogen/ThermoFisher | 404-5773-82 | 1:200 | https://www.thermofisher.com/antibody/product/FOXP3-Antibody-clone-FJK-16s-Monoclonal/404-5773-82 |
| TCR beta Monoclonal Antibody (H57-597), APC-eFluor™ 780, eBioscience™ | Invitrogen/ThermoFisher | 47-5961-82 | 1:100 | https://www.thermofisher.com/antibody/product/TCR-beta-Antibody-clone-H57-597-Monoclonal/47-5961-82 |
| Biotin anti-mouse CD185 (CXCR5), clone L138D7 | Nordic Biosite/BioLegend | 145510 | 1:100 | https://nordicbiosite.com/product/145510/Biotin-antimouse-CD185-CXCR5 |
| Pacific Blue™ anti-rat CD90/mouse CD90.1 (Thy-1.1), clone OX-7 | Nordic Biosite/BioLegend | 202522 | 1:200 | https://nordicbiosite.com/product/202522/Pacific-Bluetm-antirat-CD90mouse-CD901-Thy11 |
| PerCP/Cyanine5.5 anti-mouse CD19 | Nordic Biosite/BioLegend | 115534 | 1:200 | https://nordicbiosite.com/product/115534/PerCPCyanine55-antimouse-CD19 |
| CD279 (PD-1) Monoclonal Antibody (J43), PE-eFluor™ 610, eBioscience™ | Invitrogen/ThermoFisher | 61-9985-82 | 1:100 | https://www.thermofisher.com/antibody/product/CD279-PD-1-Antibody-clone-J43-Monoclonal/61-9985-82 |
| BV711 anti-mouse CD279 (PD-1) (clone 29F.1A12), | | | | |
| BD Horizon™ BUV737 Streptavidin | BD Biosciences | 612775 | 1:200 | https://www.bdbiosciences.com/en-dk/products/reagents/flow-cytometry-reagents/research-reagents/single-color-antibodies-ruo/buv737-streptavidin.612775 |
| BD Horizon™ BV605 Streptavidin | BD Biosciences | 563260 | 1:200 | https://www.bdbiosciences.com/en-eu/products/reagents/flow-cytometry-reagents/research-reagents/single-color-antibodies-ruo/bv605-streptavidin.563260 |
| APC Streptavidin | Nordic Biosite/BioLegend | 405207 | 1:200 | https://nordicbiosite.com/product/405207/APC-Streptavidin |
